# Extracellular Vesicles derived from Apis mellifera Royal Jelly promote wound healing by modulating inflammation and cellular responses

**DOI:** 10.1101/2022.07.21.501009

**Authors:** S. Alvarez, P. Contreras-Kallens, S. Aguayo, O. Ramirez, C. Vallejos, J. Ruiz, E. Carrasco-Gallardo, S. Troncoso-Vera, B. Morales, C.M.A.P. Schuh

## Abstract

*Apis mellifera* Royal Jelly (RJ) is a well-known remedy in traditional medicine around the world and its versatile effects range from antibacterial to anti-inflammatory properties and pro-regenerative properties. Several active compounds have been identified, however, the mechanisms of action still remain widely unknown. As a glandular product, RJ has been shown to contain a substantial number of extracellular vesicles (EVs) and in this study, we aimed to investigate the extent of involvement of RJEVs in wound healing associated effects. Molecular analysis of RJEVs verified the presence of important conserved exosomal markers such as CD63 and syntenin, as well as cargo molecules MRJP1, defensin-1 and jellein-3. RJEV internalization analysis demonstrated the involvement of membrane fusion as well as macropinocytosis or clathrin-dependent endocytosis into mammalian cells. Furthermore, RJEVs have demonstrated to modulate MSCs differentiation and secretome, as well as decrease LPS-induced inflammation in RAW 264.7 macrophages by blocking the MAPK pathway. *In vivo* studies confirmed anti-bacterial effects of RJEVs, and demonstrated an acceleration of wound healing in a splinted mouse model. Summarizing, this study suggests that RJEVs of potentially exosomal origin play a crucial role in the known effects of RJ by modulating the inflammatory phase and cellular response in wound healing.

## Introduction

Despite many advances in regenerative medicine, wound infection by bacteria remains an important problem that negatively impacts the clinical outcome of surgical procedures (1). Especially chronic wounds are prone to a number of complications such as excessive inflammation, persistent infections, formation of drug-resistant microbial biofilms, and the inability of dermal and/or epidermal cells to respond to reparative stimuli (reviewed by (2)). Over the last decades, extracellular vesicles (EVs) have become increasingly used in the field of nano-medicine. After key findings such as their role in the expression of antigens (3) and ability to transfer mRNA/miRNA (4), they are now established as one of the main factors in intercellular communication. Interestingly, it has been shown that EVs are capable of interspecies and interkingdom communication with a wide range of therapeutic applications (5).

Several honeybee products, such as honey and Royal Jelly (RJ), have been used since ancient times in various cultures for their antimicrobial and pro-regenerative properties (summarized in (6,7)). More recently, advances have been made to analyze the effects of RJ in controlled environments and on a molecular basis. Several active compounds have been found, among those Major Royal Jelly Proteins (MRJPs), Apisimin, Royalisin, Jelleins, Defensins, Apolipophorin-III-like as well as glucose oxidase (8). Pre-clinical studies described ameliorating effects of RJ in a number of conditions such as mucositis, colitis, bone formation, and infected ulcers (9–12) or diabetic foot ulcers (13). On a cellular level, RJ has been found to induce osteogenesis and exert anti-inflammatory properties in periodontal ligament cells, promote neurogenesis in neural stem cells, and also increase migration of human dermal fibroblasts (14–16). A number of studies have described the immune-modulatory properties of RJ and its components (17–19). Furthermore, antibacterial properties of RJ have first been shown in 1939 by McClesky et al demonstrating a bactericidal effect on *Escherichia coli, Eberthella thyphosa*, and *Staphylococcus aureus* (20).

Recently, our group was the first to discover the presence of extracellular vesicles as a new active part of RJ that demonstrated antibacterial and biofilm-inhibiting properties against *S. aureus* (21). Furthermore, RJ-derived EVs (RJ EVs) were found to integrate into mammalian cells and increase cellular migration in mesenchymal stem cells (21) and dermal fibroblasts (22), confirming effects previously described for crude RJ (16). The discovery that these vesicles display functional effects on mammalian cells allows hypothesizing that RJEVs might be involved in several of the known biological effects of RJ. Given their known capacity to shuttle cargo into cells and exert intracellular effects, RJEVs may serve as a potential nanomedicine-based approach for the treatment of chronic wounds in the future. Therefore, in the current interdisciplinary study, we aimed to investigate the potential of RJEVs for wound healing in a multi-model approach. We characterized RJEV cargo and modes of entry into mammalian cells to gain insight into the subsequent intracellular processing. Furthermore, we assessed the modulatory effects of RJEVs on mesenchymal stem cells and their anti-inflammatory properties as well as the signaling pathways involved. Finally, we evaluated translatability into an *in vivo* setting using a full-thickness wound mouse model.

## Materials and Methods

### EV isolation and characterization

Exosome isolation was performed as previously described (21). Briefly, RJ (Apicola del Alba, Chile) was diluted in particle-free phosphate buffered saline 1:40 (pf-PBS) and incubated for 30 min on an orbital shaker. Subsequently, debris was removed by serial centrifugation (500g-2000g, 15min each) and filtered using a 0.2µm polystyrene filter. Finally, the supernatant was ultra-centrifuged at 100.000 g for 60 min (Hanil 5 fixed rotor ultracentrifuge, Hanil, Korea) and the resulting pellet was resuspended in pf-PBS and stored at 80°C until further use.

### Nanoparticle Tracking Analysis (NTA)

As an initial step, particle quantification and size distribution were determined using NTA. RJEV samples were thawed shortly before NTA, diluted 1:100 with pf-PBS and vortexed 3 times for 1 sec. Samples were measured five times using temperature control (25°C) at camera level 8 for 60sec (NanoSight NS 3000, Malvern, UK).

### Atomic force microscopy (AFM) characterization

RJEV shapes were confirmed using AFM, as previously described (23). Briefly, freshly cleaved mica discs (12mm diameter; Electron Microscopy Sciences, US) were coated with 50µl of 0.1 M solution of poly-L-lysine (PLL, Sigma) for 5 minutes. Subsequently, discs were washed 3x with ddH2O and dried with a gentle stream of N2. RJEVs were then immobilized onto the PLL-coated mica by incubation for 30 minutes at room temperature. Samples were covered to avoid drying and potential dust contamination. After incubation, specimens were washed 3x with ddH2O and mounted onto an Asylum MFP 3D-SA AFM (Asylum Research, US). Samples were analyzed in intermittent contact mode (AC mode) with TAP300GD-G cantilevers (BudgetSensors, Bulgaria), obtaining height, amplitude, and phase channel images of substrates under environmental conditions. From the resulting height images, the surface profiles, maximum and mean vesicle height, and RMS roughness of RJEVs were calculated in the Gwyddion 2.56 software. RJEVs from three independent sample preparations were utilized for all AFM-based experiments.

### Exosomal markers and cargo

To verify exosomal origin, RJEVs were analyzed for the presence of CD63 and syntenin using Western Blot. For cargo analysis (MRJP1), RJEVs were sonicated for 15 min prior to analysis. Proteins were separated in reducing conditions (syntenin, MRJP1) and non-reducing conditions (CD63) utilizing a 10% polyacrylamide gel and blotted onto nitrocellulose. After blocking with 3% bovine serum albumin (BSA) in tris-buffered saline with 0.1% Tween (TBS-T), membranes were incubated with the respective antibodies (anti-CD63, 1:500, mouse monoclonal, Abcam ab59479; anti-syntenin, 1:1000, rabbit polyclonal, Abcam ab19903; anti-MRJP1, 1:1000, rabbit polyclonal, Cusabio CSB-PA522725EA01DNK; 3% BSA TBS-T) overnight. Subsequently, membranes were thoroughly washed and incubated with secondary antibody (IR-Dye anti-mouse, 1:15.000, LiCor; IR-Dye anti-rabbit, 1:15.000, LiCor) in 3%BSA TBS-T for 1 hour. Finally, membranes were revealed using an Odyssey CLx imaging system and results were analyzed with Image Studio Digits version 5.2.5.

Furthermore, the peptides Defensin-1 and Jellein-3 were quantified with direct ELISA, as previously described (23). Briefly, one microgram of RJEV protein and negative controls (pre-bleed serum 1:1000, human mesenchymal stem cell protein 30 μg) as well as respective standards were incubated overnight. After blocking with 5% BSA, plates were incubated with respective detection antibodies diluted in reagent diluent (R&D systems) (anti-*Apis mellifera* defensin-1, 50 ng/mL; rabbit polyclonal, GeneCust; anti-*Apis mellifera* jellein-3; 50 ng/mL, rabbit polyclonal, GeneCust) for 2 hours. HRP-conjugated anti-rabbit antibody was used at a dilution of 1:1000 (Cell signaling, UK), and developed for 10 min using substrate solution (R&D systems). The reaction was stopped with 2N H2SO4 and plates were measured using a microplate reader (Tecan Sunrise, Tecan, Austria) at 450nm.

### Effects on mesenchymal stem cells

For all experiments utilizing human mesenchymal stem cells (MSCs), written informed consent was obtained prior to MSC isolation and protocols used were approved by the Ethics Committee of Universidad del Desarrollo. MSCs derived from adipose tissue, were cultivated using standard cell culture conditions (37°C, 5% CO2) in Minimum Essential Medium (αMEM, Gibco, US) supplemented with 10% fetal bovine serum (FBS, HyClone, GE Healthcare, US), 200 mmol/L of L-glutamine (L-G) and 1% (vol/vol) penicillin-streptomycin (P/S; Sigma, US) (standard culture medium). For the experimental group “RJEV pre-conditioned MSCs”, cells were seeded at a density of 4×10^3^ per cm^2^ and incubated with 2.5×10^2^ RJEVs per cell for 48 hours. Subsequently, cells were transferred to the corresponding experiments without further addition of RJEVs.

### Routes of RJEV uptake

In order to assess routes of uptake into mammalian cells, an uptake inhibition assay was performed. RJEVs were stained with fluorescent dye CFSE (Cell Trace CFSE, Thermo Fisher Scientific), as previously described (24). Human adipose-derived mesenchymal stem cells were seeded at a density of 5.000 cells per cm^2^ in a 6-well plate and left to adhere overnight. Each potential uptake route was blocked with either the inhibitor Omeprazole 1 mM, Amiloride 50 µM, Chlorpromazine 28 µM, or Filipin 7.5 µM (for 15 min). Cells were subsequently incubated with 2.5×10^3^ CFSE-RJEVs per cell for 4 hours and analyzed with flow cytometry (CyAn ADP, Beckman Coulter, US). Treatment groups were compared to unstained control as well as CFSE-RJ-EVs without inhibitor. For data analysis, the FlowJo software (TreeStar, version 8.8.6) was used.

#### Population Doubling Time

To assess the effects of RJEVs on population doubling time, MSCs were seeded at an exact density (3.3×10^3^ per cm^2^) and left to adhere for 1 hour, followed by the addition of 2.5×10^2^ RJEVs per cell at every passage (RJEV group). Cells were counted after 120 hours and reseeded at the same density until passage 13. Calculation was done with the following formula: PDT in days = ((Time in days) * log (2)) / (log (final cell number) * log (initial cell number)).

#### Secretory protein profile

To assess changes on the secretory profile of MSCs in presence of RJEVs, four proteins involved in wound healing were selected for analysis: fibroblast growth factor (FGF), vascular endothelial growth factor (VEGF), insulin-like growth factor (IGF-1a), and hepatocyte growth factor (HGF). Briefly, 10^3^ cells per cm^2^ were seeded in a 6-well plate and left to adhere overnight. For conditioning, cells were cultivated in media without FBS. Supernatants were collected after 24h, centrifuged at 1.000xg for 15 min to remove cellular debris, and stored at -80°C until analysis. The respective proteins were quantified with sandwich ELISA (DuoSet, R&D Systems with ancillary kit) according to manufacturer’s instructions. Finally, the results were measured using a microplate reader (Tecan sunrise, Tecan, Austria) at 450nm.

#### Differentiation into adipogenic, osteogenic and chondrogenic lineage

Cells were seeded at the respective cell densities for each differentiation assay (adipogenic lineage: 7.5×10^3^/cm^2^; osteogenic lineage: 5×10^2^/cm^2^; chondrogenic lineage: 7.5×10^3^/cm^2^; undifferentiated control 5×10^2^/cm^2^) and were left to adhere overnight in standard culture medium. Subsequently, cells were incubated with respective differentiation media (StemPro adipogenic, osteogenic, chondrogenic differentiation kits; Gibco, US) for 14 days. An undifferentiated control was cultivated in standard culture medium supplemented with 5% FBS. RJEVs (10^6^/well) were added and each medium was changed every 3 days for 14 days. Adipogenic and osteogenic differentiation were evaluated using Oilred O and Alizarin Red staining, as previously described (25). For chondrogenic differentiation, cells were fixed with ice-cold 70% methanol for 10 minutes at room temperature, and subsequently stained with 0.1% Safranin O (w/v) for 10min. For quantification, 200µl 100% ethanol were added to each well and the plate was incubated for 30 min at an orbital shaker. Prior to quantification, representative images were taken on a Nikon Eclipse TS100. Quantification was performed using a microplate reader (Tecan sunrise, Tecan, Austria)

### Anti-inflammatory assays

The anti-inflammatory potential of RJEVs was assessed by employing the RAW 264.7 macrophage cell line. RAW264 cells were cultivated using standard cell culture conditions (37°C, 5% CO2) using certified LPS-free Dulbecco’s Modified Eagle Medium and supplements (10% FBS, 200 mmol/L L-G, 1% P/S). For all anti-inflammatory assays, cells were seeded at a density of 2×10^3^ cells/cm^2^. RJEVs were added at concentrations of 250/cell and 2.5×10^3^/cell. For determining the secretory profile of pro-inflammatory cytokines (Interleukin 1 β (IL1β), IL6 and TNFα), RAW 264 cells were incubated with RJEVs for 12 hours, and subsequently stimulated with 1µM LPS for 24h. Supernatants were collected, centrifuged at 1.000xg for 15 min to remove cellular debris, and stored at -80°C until analysis. Respective proteins were quantified with sandwich ELISA (DuoSet, R&D Systems with ancillary kit), according to manufacturer’s instructions. Results were measured using a microplate reader (Tecan sunrise, Tecan, Austria) at 450nm. For analysis of the associated pathways, RAW 264 cells were incubated with RJEVs for 12 hours, and subsequently stimulated with 1µM LPS for 6h. Cells were lysed using RIPA buffer for 15 min on ice, centrifuged at 10.000g to remove cellular debris, transferred to another tube, and stored at -20°C until further analysis. For Western Blot, proteins were quantified using BCA assay. Proteins were separated in a 10% polyacrylamide gel, blotted onto nitrocellulose membranes, and blocked with 3% BSA-TBS-T. Membranes were incubated with the phosphorylated antibodies prior to unphosphorylated. Membranes were then thoroughly washed and incubated with secondary antibody (IR-Dye anti-mouse, 1:15.000, LiCor; IR-Dye anti-rabbit, 1:15.000, LiCor) in 3% BSA TBS-T for 1 hour. Membranes were revealed using an Odyssey CLx imaging system, and results were analyzed with Image Studio Digits version 5.2.5.

### Effects of RJEVs in vivo

All experimental procedures were approved by the local animal committee in accordance with the Chilean law and ARRIVE guidelines as well as 3R practices. C57BL/6 mice were kept in an enriched environment with a 12h/12h light/dark cycle, at 23°C-25°C ambient temperature and food as well as water *ad libitum*. For experimentation, 10– 12-week-old male and female mice with a minimum weight of 20g were utilized. Well-being of animals was assessed daily using a Grimace scale as well as Morton and Griffith assessment. In all *in vivo* experiments, RJEVs were administered using a collagen type I hydrogel as previously described (22). Briefly, 24h prior to experimentation, collagen from rat tail (2 mg/ml in hydrochloric acid; Gibco, US) was mixed with 10xMEM (Gibco, US) in a ratio of 8:1. Gelation was induced by neutralization with 1M Sodium Hydroxide and 1 part RJEVs were added at concentration of 5×10^9^/ml. Control gels not carrying RJEVs were adjusted with 1 part PBS. The resulting solution was transferred into a 96-well plate (100µl), and incubated at 37°C for 30 min. Subsequently, gels were covered with 100µl PBS.

#### In vivo antibacterial properties

Prior to surgery, *Staphylococcus aureus* ATCC 25923 cells were activated in BHI broth at 37°C and 150rpm orbital shaking for 5 hours and adjusted to 0.5 Mc Farland directly before experiments. Mice were randomly assigned to respective groups (ctrl, *S. aureus* ctrl, collagen ctrl, collagen RJEV) and deeply anesthetized using sevoflurane by inhalation. After hair removal on both sides of the dorsum, the area was disinfected with povidone-iodine in order to avoid contamination. A 5 mm incision was made on both sides of the dorsum and undermining was performed to create a small pocket. Collagen gels (collagen ctrl, collagen RJEV) were cut in half to create a semilunar shape (2.5×4×1mm), and placed inside the pocket. To the ctrl, collagen ctrl, and collagen RJEV groups, 5µl of bacterial solution were added. SHAM received 5µl of saline solution. Surgical sites were closed with 5.0 vicryl sutures. Mice were given analgesics (Meloxicam, 2mg/kg) after the surgery and the following day. 48h after surgery, mice were euthanized and incision sites were opened. Using a sterile loop, bacteria were collected from the pocket and placed in 1 ml BHI broth. Subsequently, bacteria were plated in serial dilution on BHI agar plates for colony counts, and in 96-well plates (100µl BHI broth) for viability assays. After over-night incubation, colonies were counted on agar plates. For viability quantification, resazurin (0.016%, Sigma-Aldrich) was added to the wells for 1h at 37°C, and was subsequently measured at 570nm in a microplate reader.

#### Mouse Wound Healing Model

Mice were randomly assigned to respective groups (ctrl, collagen ctrl, collagen RJEV) and deeply anesthetized using sevoflurane by inhalation. The center part of the dorsum was depilated around the neck, and disinfected with povidone-iodine. Subsequently, a 5 mm full-thickness wound was created using a sterile histopunch. A silicone wound splint (15 mm/7 mm outer/inner diameter, Grace Biolabs, UK) was centered on the wound area and fixed with skin adhesive as well as 6 sutures (vicryl 5.0). Collagen gels were placed on the fresh wound and the area was covered with Tegaderm. Mice were given analgesics (Meloxicam, 2mg/kg) once a day for the three days after surgery. Collagen gels were replaced every 5 days and the wound area was covered again with Tegaderm. Pictures were taken directly after surgery, as well as on day 5, 10 and 15. Wound closure was measured using ImageJ software and calculated in comparison to day 0. On day 15, mice were euthanized, and tissue was harvested and fixed (4% formaldehyde) for 48h. For histological analysis, tissues were embedded in paraffin using standard protocols. 5-µm sections were stained with Hematoxylin/Eosin to assess re-epithelization, vascularization and inflammation using a score system, as described by de Moura Estevao et al (26).

### Statistics

All data in this study are shown as mean ± standard deviation. All data were tested for normality using Shapiro-Wilk test. Statistical analysis was performed using student-t test (for two groups) or one-way ANOVA followed by Tukey’s range test for significant differences between means (for three or more groups). For time-course analysis, two-way ANOVA with Šídák’s multiple comparisons test was used. Analysis was performed using GraphPad Prism 5 for Mac OS X, version 9.3.1 (GraphPad Software, Inc.). Significance was considered at p<0.05 (see figure legends for specific values). N values were determined from independent experiments and independent isolations of RJEVs.

## Results

### RJEV characterization and uptake analysis

Prior to functional experiments, we performed an in-depth characterization of RJEVs utilized in this study. Isolated RJEVs were found to display a mean size of 134.5 +/- 16.5 nm and a median size of 124.2 +/- 14.9 nm, demonstrating RJEV homogeneity within the samples. No significant differences were found between the mean and median RJEV sizes. Particle yield was between 3.9×10^9^ and 2.1×10^9^ particles per gram raw material, which is in line with previous observations (Fig. 1A) (21). Furthermore, we utilized AFM to characterize the ultrastructure of mica-immobilized RJEVs, confirming vesicle shape and integrity at the nanoscale in both height and amplitude images (Figure 1B). Also, RJEVs were found to display a maximum height (17.6 +/- 4.9 nm) and mean height (9.7 +/- 4.2 nm) consistent with the expected morphology of mica-bound EVs, and an average RMS surface roughness of 5.4 +/- 1.2 nm.

**Figure 1:**
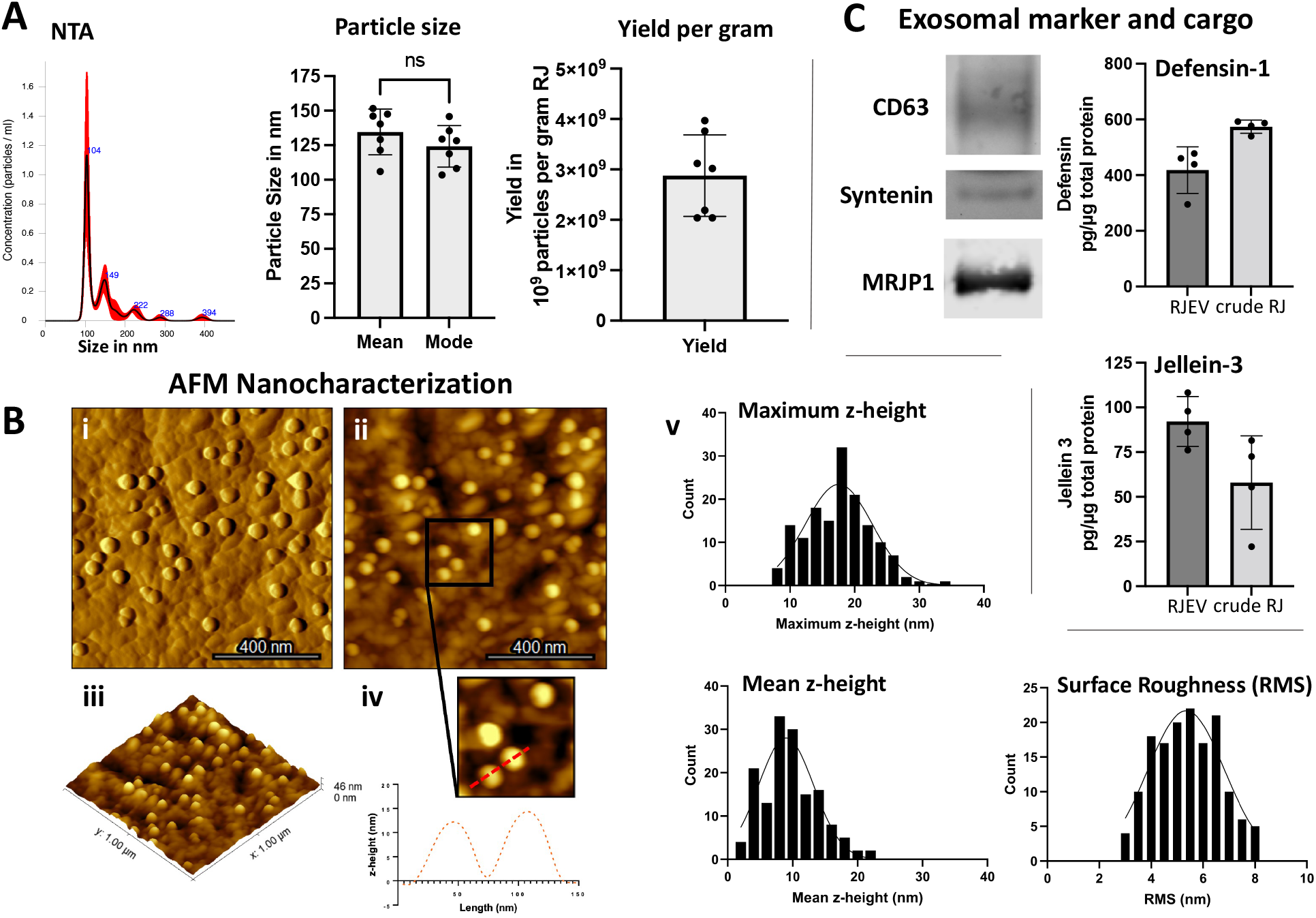
Morphological and biochemical characterization of RJEVs; **A)** representative NTA histogram; average (mean) and median (mode) particle size in nm (*p<0.05; student t-test); Particle yield of RJEV per gram raw RJ; n=7; **B)** AFM-based analysis of RJEV vesicle shape; **i)** amplitude, **ii)** height, and **iii)** 3D height reconstruction of RJEV immobilized onto PLL-coated mica. iv) Profile trace obtained on 2 RJEV demonstrating their shape and size. **v)** Maximum and mean heights and RMS surface roughness of immobilized RJEV (n=150). **C)** Western Blot analysis of exosomal markers. CD63, Syntenin-1 and RJEV cargo MRJP-1; n=3; ELISA analysis of RJEV cargo Defensin-1 and Jellein-3; n=4;

Exosomal origin of RJEVs was confirmed by verifying the presence of CD63 and Syntenin, two conserved exosomal markers present in *Apis mellifera* (KEGG: exosome: *Apis mellifera*; entry 552721 CD63; entry 551650 syntenin). Furthermore, the relevant cargo proteins MRJP-1, Defensin-1 and Jellein-3 were also identified within RJEVs.

In order to study uptake mechanisms, the specific inhibitors amiloride, omeprazole, filipin III, and chlorpromazine were used to assess the role of macropinocytosis, membrane fusion, lipid raft or caveolae-mediated endocytosis, and clathrin-dependent endocytosis, respectively. Amiloride, omeprazole, and chlorpromazine significantly decreased uptake of CFSE-RJEVs into MSCs, to similar extents (64% +/- 12.15%, 65.9% +/- 11.6%, 64.51% +/- 11.2%). However, filipin III did not exert significant inhibitory effects on CFSE-RJEV uptake (94.9% +/- 8.8%), indicating a minor role of lipid rafts in RJEV internalization (Fig. 2).

**Figure 2:**
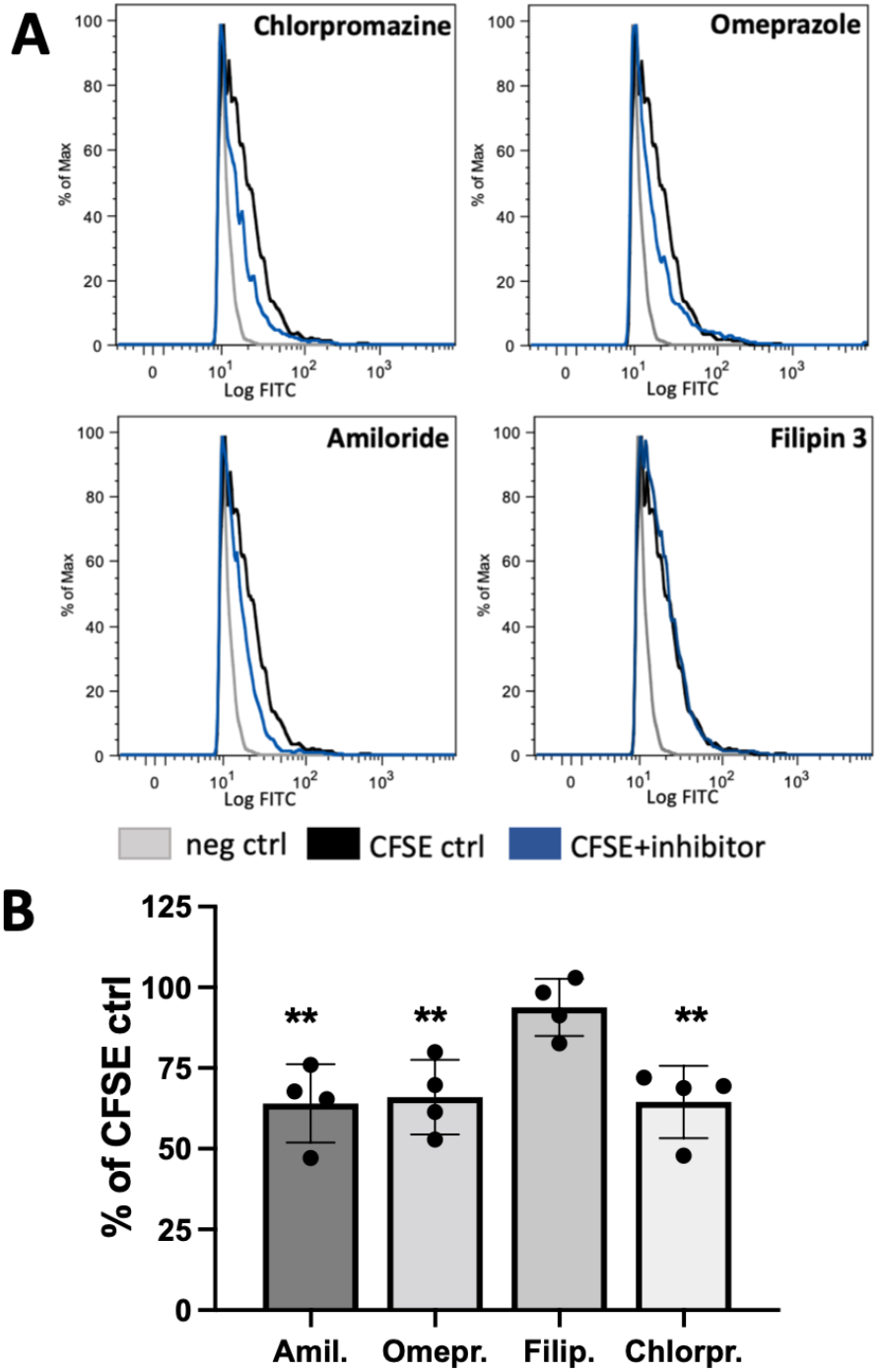
Route of RJEV uptake into human MSCs; **A)** Flow cytometry histograms of MSCs without RJEV (neg ctrl, grey), incubated with CFSE-RJEV (CFSA ctrl, black) and with CFSE-RJEV and the respective inhibitor (blue, CFSE+inhibitor) chlorpromazine, omeprazole, amiloride and filipin 3; **B)** quantitative analysis of RJEV uptake inhibition into MSCs, displayed as percentage of respective CFSE-RJEV control; n=4, (*p<0.05; **p<0.01; one-way ANOVA with Tukey’s post-hoc);

### Effect of RJEVs on MSC differentiation, population doubling time, and secretion

As a next step, MSC cultures derived from adipose tissue of healthy donors were analyzed for in vitro chondrogenic, osteogenic, and adipogenic differentiation potential, either pretreated or in the presence of RJEVs (Fig. 3A/B). Chondrogenic differentiation was confirmed with Safranin O staining, and both qualitative and quantitative analysis showed a significant increase (25-40%) of proteoglycan deposition in the RJEV group, compared to control; however, no effect was observed in the pre-treatment group. Interestingly, alizarin red stain for calcium after osteogenic induction was increased in both groups, with RJEV displaying a highly reproducible effect of around 50% increase compared to the untreated control. Pre-treatment resulted in an increase of 10%-130% compared to control. Furthermore, adipogenic differentiation analyzed by Oilred O was significantly decreased in the RJEV group compared to untreated control. Pre-differentiation, comparable to osteogenic differentiation, lead to a higher spread within the data and a less predictable outcome. While three out of four donors displayed a decrease in adipogenic differentiation, one donor in the pre-treatment group displayed an increase (Fig. 3A/B).

**Figure 3:**
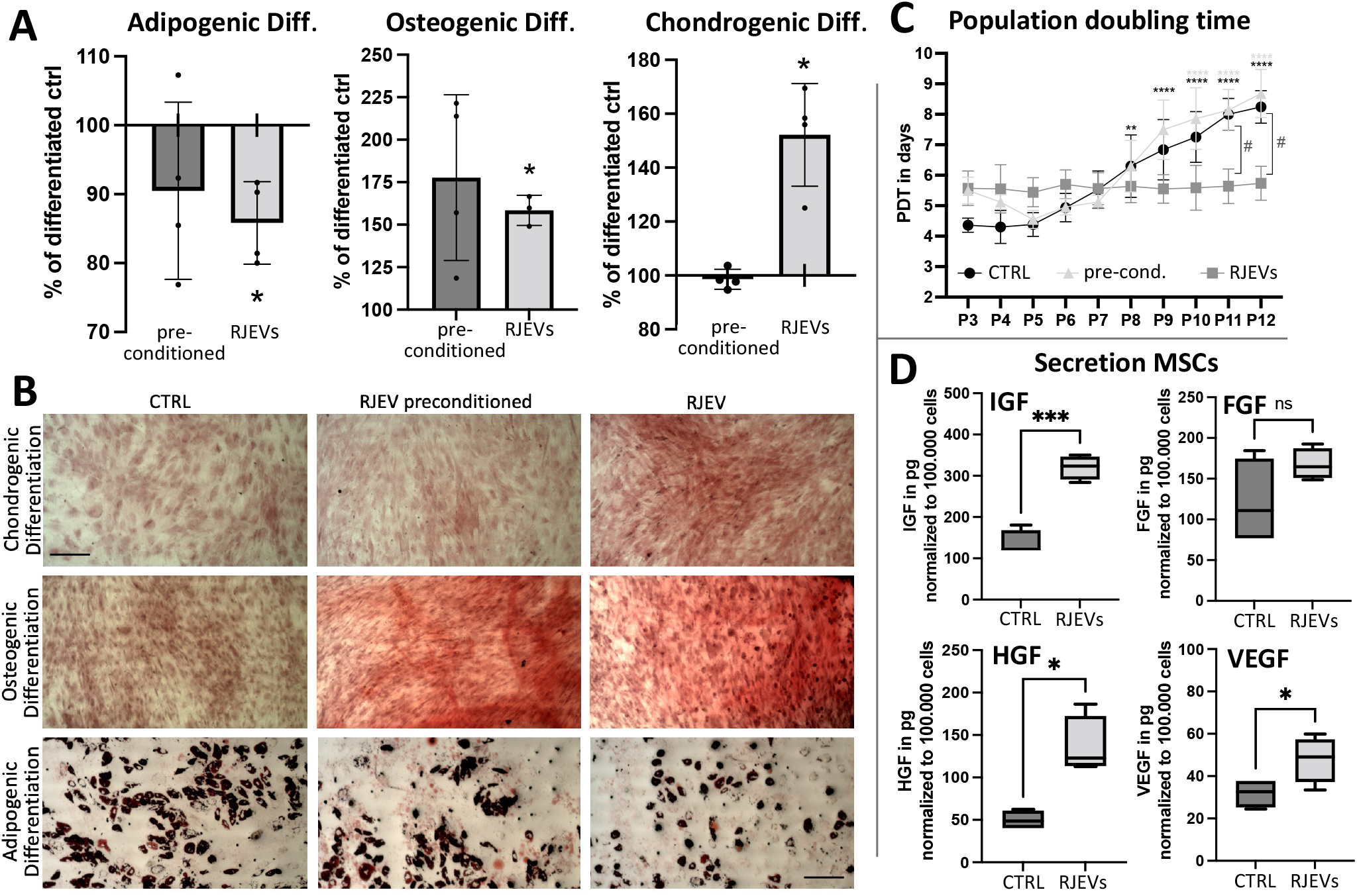
Effects of RJEVs on human MSCs; A/B) Differentiation into adipogenic, osteogenic and chondrogenic lineage; A) quantification of differentiation in RJEV preconditioned group (dark gray) and RJEV group (light gray), displayed as percentage of untreated differentiated ctrl (≙ 100%); n=4, with technical duplicates; statistical significance tested with one-way ANOVA with Tukey’s post-hoc; *p<0.05; **B)** Representative images of MSC differentiation after 14-day induction and with Oilred O (adipocytes), Alizarin Red (osteocytes) and Safranin O (chondrocytes); scale bar ≙ 100µm; **C)** population doubling time of MSCs; seeded at 3.3×10^3^ per cm^2^, passaged and counted every 120h (5 days); calculation: PDT in days=((Time in days) * log(2)) / (log (final cell number) * log (initial cell number)); groups: untreated ctrl (black, circle), RJEV preconditioned MSCs (light gray, triangle) and RJEV (dark gray, square); Significance tested with two-way ANOVA and Šídák’s multiple comparisons test; *p<0.05; **p<0.01, ***p<0.001, ****p<0.0001, of respective group in comparison to P3; # p<0.05, between groups within the same passage. D) Secretory profile of MSCs in presence of RJEV; levels of IGF-1, FGF, HGF and VEGF in supernatants measured with ELISA; secretion normalized to 100.000 cells. N=4; significance tested with students t-test; *p<0.05; **p<0.01, ***p<0.001;

Subsequently, population doubling time (PDT) was observed over a time period of 10 passages (p3-p12) (Fig. 3C). In the control group, a continuous increase in PDT after P3 was observed. In P8-P12, PDT was significantly higher compared to P3. In the RJEV group, while initial PDT was higher in P3-P6, the PDT remained approximately the same over the course of 10 passages (5.6 +/- 0.5 days, in comparison to control 6 +/- 1.6 days and RJEV preconditioning 6.37 +/- 1.7 days). Interestingly, MSCs preconditioned with RJEVs displayed a PDT similar to RJEVs in P3, subsequently decreased in P4 and P5, and finally increased to levels comparable with the control group, indicating a transitory effect of RJEV preconditioning on proliferation.

Aside from the potential of MSCs to differentiate into specific lineages, their secretory profile has also been shown to be highly important (Fig. 3D). Therefore, we assessed the influence of RJEVs on the secretion of four factors known to be involved in wound healing, and found that IGF (133.65+/- 32.63 ng to 320.36 +/- 28.77 ng/10^5^cells), HGF (49.9 +/- 11.21 ng to 136.22 +/- 34.34 ng/10^5^cells), and VEGF (31.86 +/- 6.8 ng to 47.87 +/- 10.88 ng/10^5^cells) were significantly increased in the RJEV group. Finally, there also was a tendency towards a higher secretion of bFGF in the presence of RJEVs, however, the results were not statistically significant.

#### Anti-inflammatory effect of RJEVs

Royal Jelly has been previously described as anti-inflammatory; however, so far it is unknown how RJEVs are involved in this effect. Hence, we stimulated RAW 264.7 macrophages with LPS and subsequently analyzed the secretion of the pro-inflammatory cytokines IL1β, IL6, and TNFα. RJEVs in absence of LPS did not trigger increased secretion of pro-inflammatory cytokines, indicating that RJEVs alone are not pro-inflammatory (Fig. 4A). LPS significantly increased the secretion of all three pro-inflammatory cytokines assessed. Presence of higher dose RJEVs (10^7^ RJEVs) during LPS stimulation significantly decreased the amount of secreted IL1β, IL6, and TNFα (56.32%, 58.08% and 63.04% decrease compared to LPS control, respectively). Lower dose (10^5^ RJEVs) displayed significant effects in IL6 and TNFα secretion (15.99% and 24.58% decrease), however, not in secretion of IL1β (9.8%). (Fig. 4A)

**Figure 4:**
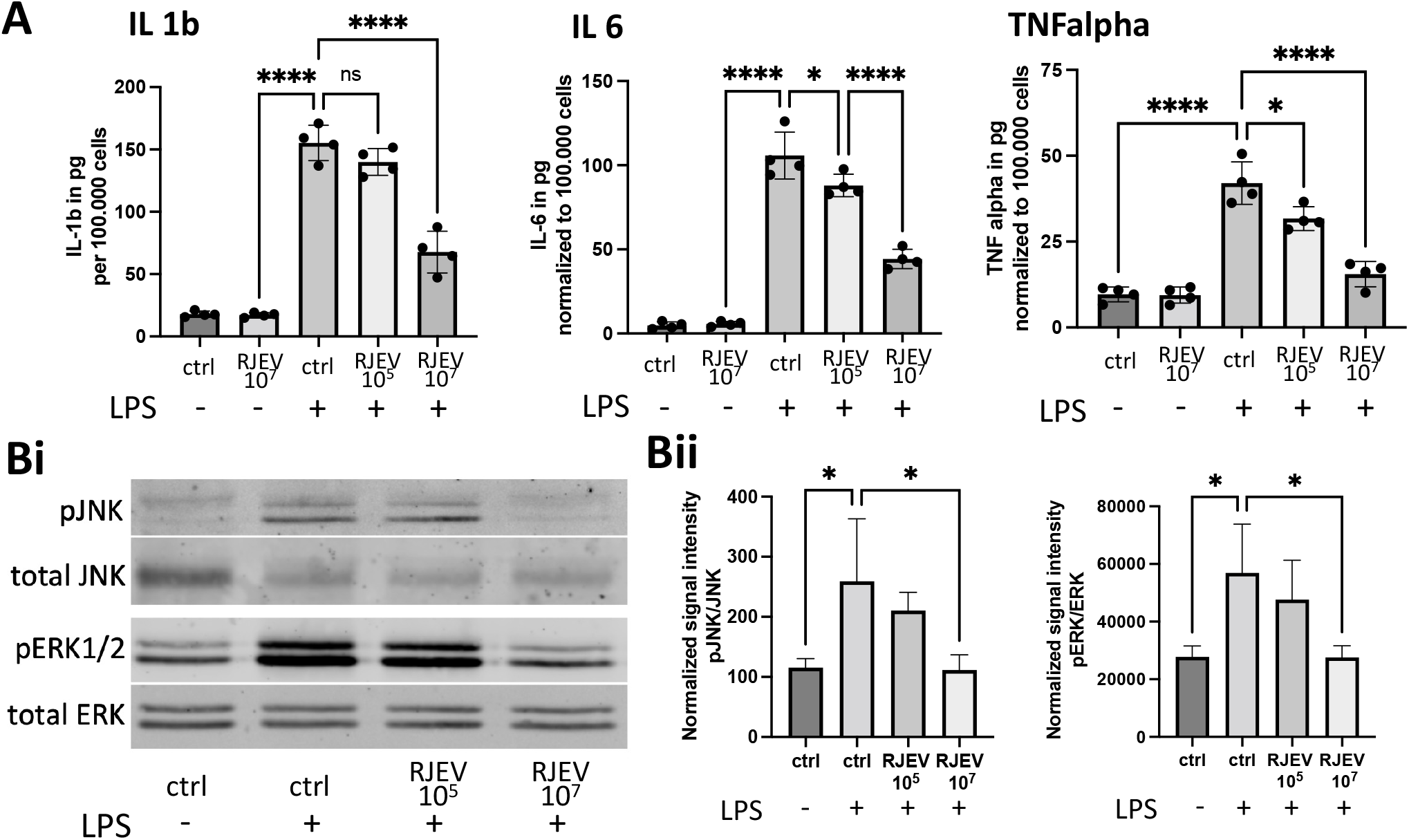
Anti-inflammatory effect of RJEVs; **A)** secretion of pro-inflammatory cytokines IL-1, IL-6, TNFα from RAW 264.7 macrophages after stimulation with LPS, measured with ELISA; secretion was measured in cells without LPS stimulation and treatment (ctrl, LPS-), in presence of RJEV (RJEV 10^7^, LPS-), stimulated with LPS (ctrl, LPS+), with low dose of RJEV (RJEV 10^5^, LPS+), and high dose of RJEV (RJEV 10^7^, LPS+); results were normalized on 100.000 cells; n=4; **Bi)** Western Blot analysis of phosphoERK/ERK 1/2, and phosphoJNK/JNK expression in RAW 264.7 macrophages of treated with RJEV 10^5^ and RJEV 10^7^, stimulated with LPS, compared to untreated and unstimulated ctrl, visualized with Odyssey; **Bii)** Quantification of phosphoERK 1/2 relative to ERK 1/2 and phosphoJNK relative to JNK in respective groups; n=4; Significance for A) and C) was tested with one-way ANOVA with Tukey’s post-hoc; *p<0.05, **p<0.01, ***p<0.001, ****p<0.0001;

Finally, Western Blot analysis of JNK and ERK1/2 revealed a strong response of both to LPS, expressed in phosphorylation of both proteins. In a dose dependent manner, RJEVs significantly decreased phosphorylation, indicating an involvement of RJEVs in the MAPK pathway (Fig. 4Bi/Bii).

#### In vivo antibacterial properties

The antibacterial properties of RJEVs were demonstrated by our group previously in several studies under controlled *in vitro* conditions; however, this effect had not yet been confirmed in an *in vivo* model. In order to create controlled conditions, a subcutaneous pouch model was chosen to avoid cross-contamination with other bacterial strains, wound cleaning, and systemic infection, among others. Colony count after 48h *in vivo* and subsequent 24h cultivation on BHI agar, revealed around 8.0×10^7^ CFU in the *S. aureus* control group and 7.62×10^7^ CFU in the *S. aureus* collagen gel group (Fig. 5B). Most importantly, no bacteria were detected in the control group without *S. aureus* and the *S. aureus* RJEVs collagen gel group. This result was confirmed by viability tests in a microwell plate and resazurin, demonstrating a bactericidal effect of RJEVs released by collagen gels *in vivo (Fig. 5C)*.

**Figure 5:**
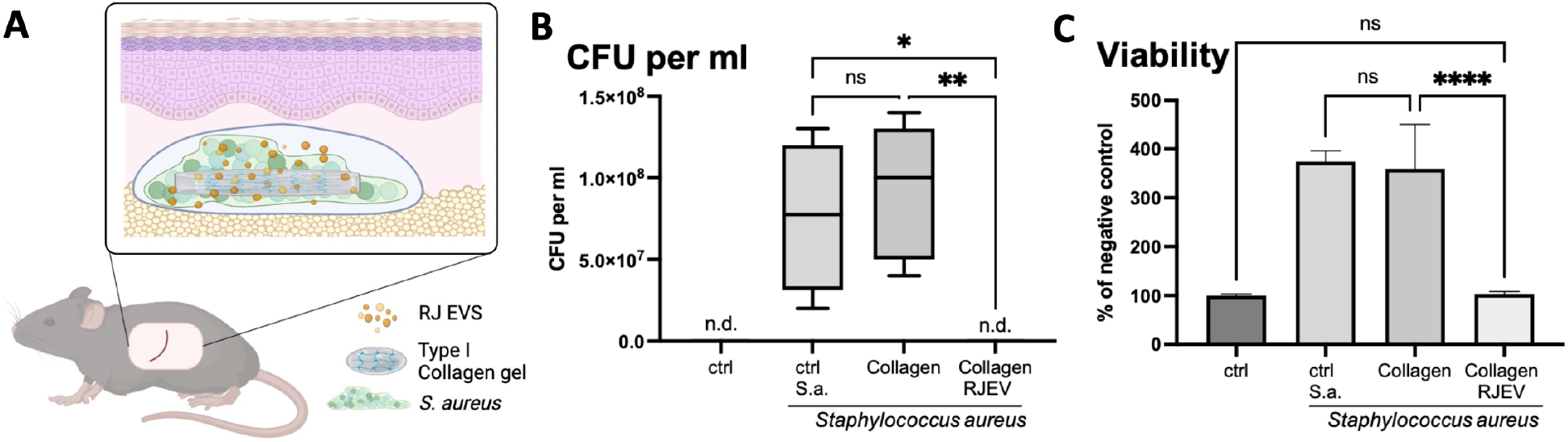
Antibacterial activity of RJEVs in vivo; **A)** schematic of animal model utilized to assess antibacterial activity; in 10-12 week old C57BL6 mice a subcutaneous pocket was created and *S. aureus* ATCC 25923 were incubated in with type I collagen gels containing 5×10^9^ RJEV/ml, compared to SHAM ctrl, untreated ctrl and collagen gel ctrl without RJEVs. The collagen gel releases RJEV over time, which are exerting an antibacterial effect, evaluated after 48h in vivo; **B)** Quantification of colony forming units (CFU) per ml; n.d. not detected and **C)** viability measured with resazurin assay in SHAM, *S. aureus* ctrl, collagen, and collagen RJEV; n=4-6; Significance for B) and C) was tested with one-way ANOVA with Tukey’s post-hoc; *p<0.05, **p<0.01, ***p<0.001, ****p<0.0001.

#### In vivo wound healing assay

Finally, the effect of RJEVs on epithelial regeneration was assessed in a full-thickness wound mouse model. After infliction, wounds were covered with collagen hydrogels with and without RJEVs and wound closure was documented every 5 days. RJEVs significantly accelerated wound healing in the first 10 days, compared to collagen and untreated control (Fig. 5Bi/Bii); however, on day 15 all wounds were completely closed, and no difference could be observed on a macroscopic level. Furthermore, no important differences in vascularization were observed between the groups. Nevertheless, inflammatory cell and epithelization quantification showed that RJEVs significantly improved wound healing on a cellular level (Fig. 6Ci/Cii).

**Figure 6:**
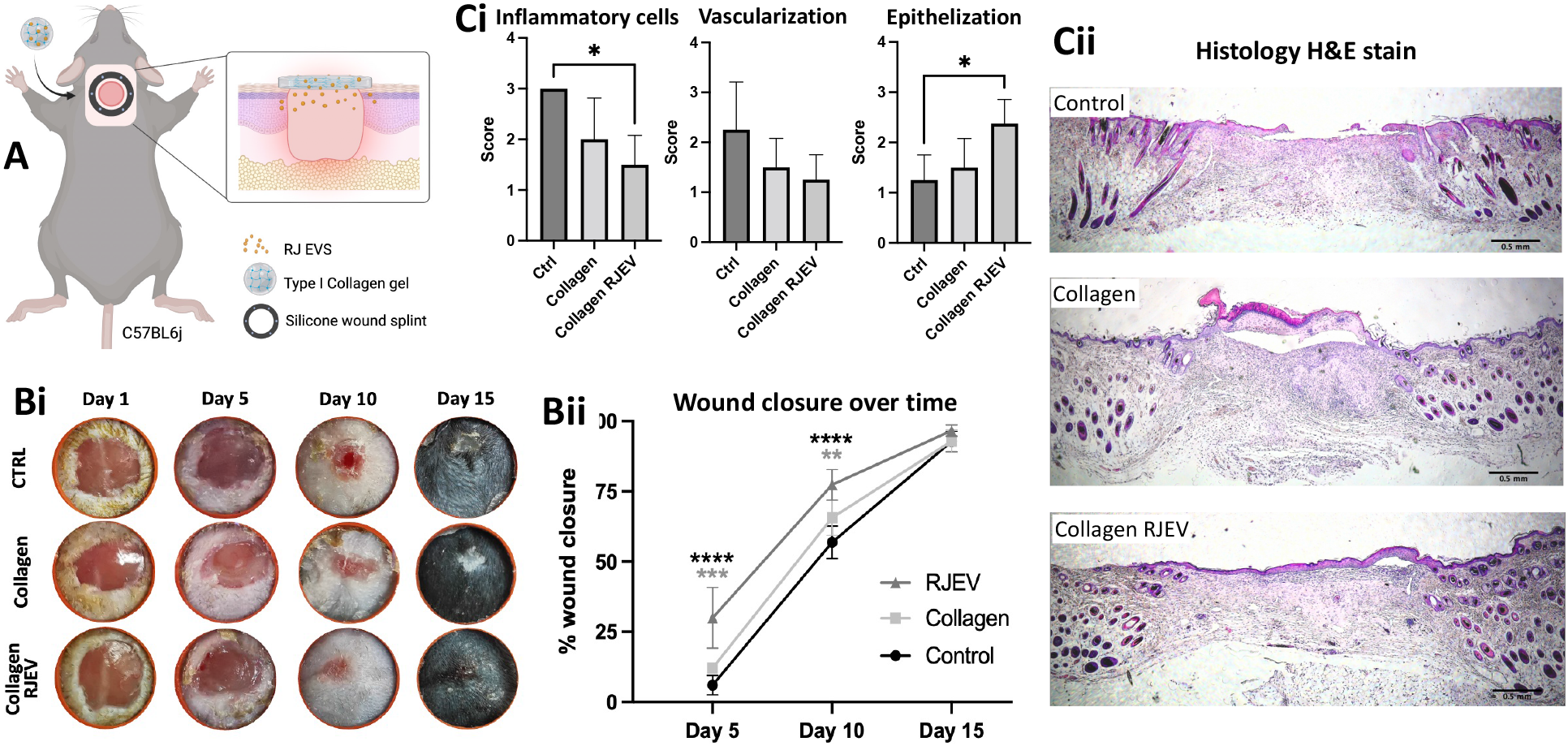
Effects of RJEVs in a splinted mouse wound healing model; **A)** schematic of the wound healing model utilized; a full-thickness wound was inflicted in 10-12 week C57BL6 mice, and splinted with a silicone ring to avoid contraction; wounds were treated with type I collagen gels releasing RJEVs; **Bi)** Representative images of silicone splinted excision wounds at day 0, 5, 10 and 15; **Bii)** quantification of wound closure over time, displayed in percentage compared to day 0; n=7; significance was tested with one-way ANOVA with Tukey’s post-hoc for differences between groups on different timepoints; *p<0.05, **p<0.01, ***p<0.001, ****p<0.0001; black asterisk displays difference to ctrl, gray asterisk to collagen group; **Ci)** histoscore quantification for inflammatory cells, vascularization and epithelization; n=4 significance was tested with one-way ANOVA with Tukey’s post-hoc for differences between groups on different timepoints; *p<0.05, **Cii)** representative images of the wound tissues harvested on day 15, stained with hematoxylin/eosin; scale bar ≙500 µm;

**Figure 7:**
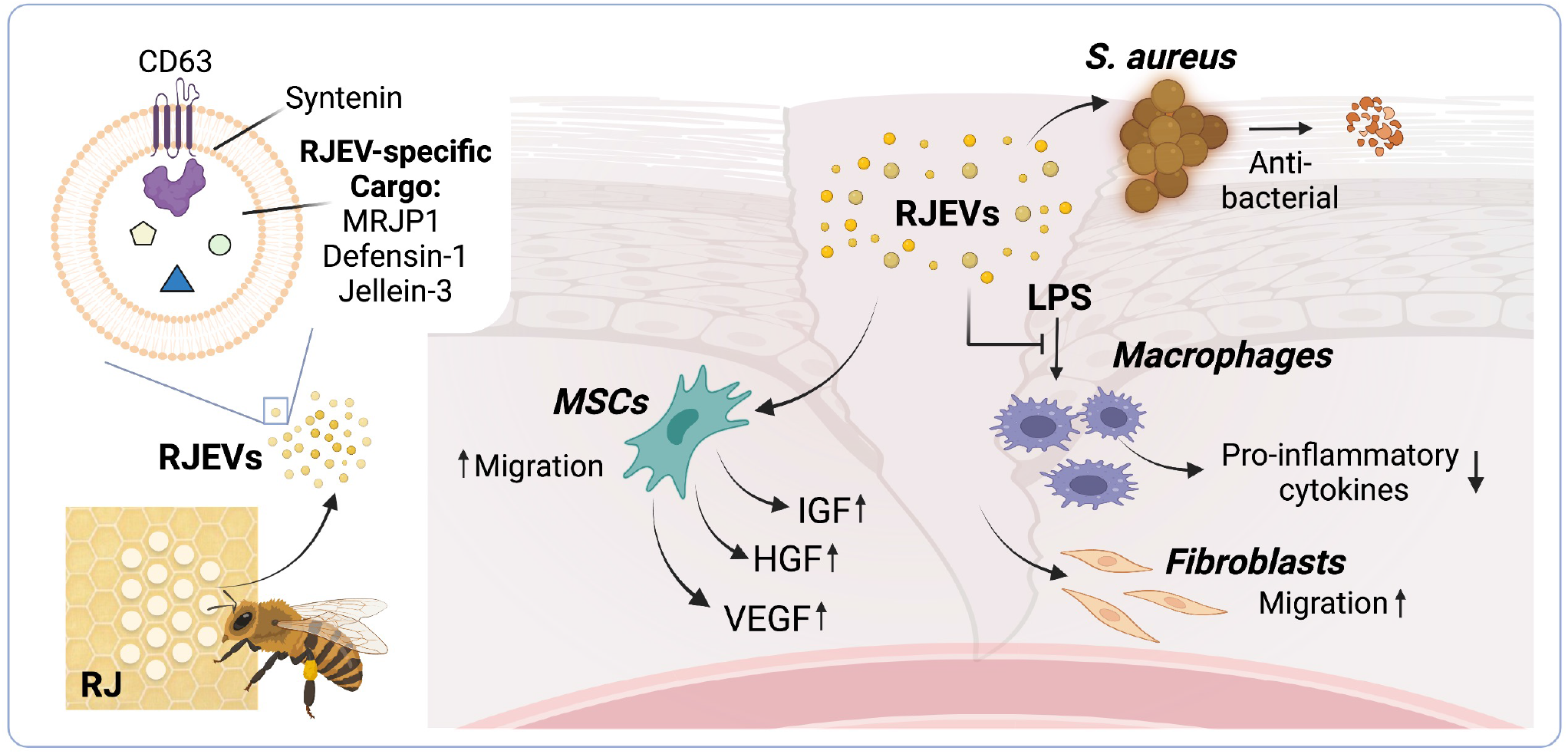
Graphical summary of RJEV findings; isolated RJEVs display markers associated with exosome biogenesis (CD63, syntenin) and contain RJ-specific cargo proteins MRJP1, defensin-1, and Jellein-3; RJEVs are antibacterial against *S. aureus in vitro* (21,22) and *in vivo*; in a full-thickness wound mouse model, RJEVs accelerated wound closure and improved epithelization. The underlying effects may be associated with the modulation of cells involved in the wound healing process, including fibroblasts immune cells such as macrophages, as well as tissue-resident MSCs. Fibroblasts and MSCs have shown an increased migratory behavior in presence of RJEV (21,22), and MSCs display changes in the secretome, increasing the secretion of pro-regenerative factors IGF, HGF, and VEGF. In macrophages, RJEVs blocked the MAPK pathway activated by LPS, resulting in a decreased secretion of pro-inflammatory cytokines IL1β, IL6, and TNFα.

## Discussion

Extracellular vesicles derived from non-mammalian sources have become increasingly interesting in the last decade as a potential tool for the standardization of natural medical treatments (5). However, the characterization of novel non-mammalian EV sources and EVs has been proven complex due to the absence of classical exosomal markers and the need to find validated alternatives, workarounds, and novel criteria. Honeybee EVs have first been reported by our group in a preliminary study that assessed vesicle shape and internalization into mammalian cells (21). Particle characterization and quantification for this study revealed a relatively pure EV population, with no significant secondary populations (Fig. 1A). Verification of vesicle shape was previously performed using transmission electron microscopy (21). However, while this method displays a number of advantages, such as broad availability and easy-to-use protocols, it lacks further insight into particle characteristics such as height and surface structure. Furthermore, electron microscopy involves destructive sample preparation (e.g. counterstaining, vacuum) that can alter the ultrastructural morphology and properties of EVs. Therefore, here we utilized AFM in tapping mode in order to gain information on RJEV shape and ultrastructural morphology with minimal sample destruction under environmental conditions. At the nanoscale, RJEVs were found to display similar sizes, shapes, and morphological characteristics as other EVs and exosomes from diverse sources previously described in the literature (27–29). Overall, AFM quantitative analysis showed an average maximum height of 17.6 +/- 4.9 nm and a mean height of 9.7 +/- 4.2 nm for RJEVs. Furthermore, a homogenous nanoscale surface roughness was observed across all studied RJEVs (mostly ranging between 4-7 nm). As vesicle surface roughness is mostly determined by membrane composition (30), particle roughness analysis may serve as a reference value for future EV studies utilizing mammalian or non-mammalian sources.

In order to determine the origin of EVs, classical cytosomal and membrane markers associated to the endosomal sorting complex required for transport (ESCRT) machinery are commonly analyzed. However, both the ESCRT-dependent as well as ESCRT-independent pathways have been mainly characterized for mammalian cells (31), while for e.g. insects and plants mechanisms have yet to be elucidated. Therefore, we assessed the potential exosomal origin of RJEVs by employing the *Kyoto Encyclopedia of Genes and Genomes* (KEGG) to identify candidate proteins for *Apis mellifera*. CD63 and syntenin (KEGG: exosome: *Apis mellifera*; entry 552721 CD63; entry 551650 syntenin), which are crucial for ESCRT machinery were both found in the RJEV samples (Fig. 1C). Syntenin is known to recruit CD63 in early endosome formation (32,33), indicating a potential exosomal origin of the isolated RJEVs. Nevertheless, further studies are needed to identify the exosome biogenesis and release mechanism in honeybees, since several well-known components have been described in *Apis mellifera*, such as ESCRT-I, II, or Syndecan (KEGG: *Apis mellifera*; entry 551408, 410869, 413557), but several others such as ESCRT-III or ALIX are missing.

As mentioned above, our previous studies reported the uptake of RJEVs into mammalian cells, which we further investigated in this study. We hypothesized an uptake of RJEVs dependent on macropinocytosis or endocytosis, since the differences in membrane composition between mammalian cells and honeybee cells are significant in terms of composition (e.g. presence of bee-specific fatty acids such as 10-HDA (34)), and may not facilitate membrane fusion. Interestingly, we found membrane fusion as well as macropinocytosis and clathrin-dependent endocytosis to be involved in RJEV uptake into mammalian cells (Fig. 2). This indicates a dual effect of RJEVs in mammalian cells: on the one hand, a direct release of RJEV cargo (e.g. MRJP-1, Defensin-1, Jellein-3, Fig. 1C) into the cytoplasm after membrane fusion; and on the other hand, intracellular sorting following internalization via other mechanisms. Interestingly, lipid raft-mediated endocytosis (blocked by Filipin-III) appeared not to be part of the RJEV uptake mechanism (Fig. 2), which might be explained by missing interactions of honeybee-specific fatty acids as well as other membrane components with mammalian cell lipid rafts.

Royal Jelly has a longstanding history of medical applications, especially in accelerating wound healing (9–12,35), as well as displaying anti-microbial (36–41) and anti-inflammatory effects (17). The underlying effects have been attributed to a number of effects, including Royalactin, also known as MRJP-1, contributing to increasing multipotency of mammalian stem cells (42). Since we could verify the presence of MRJP-1 within RJEVs, we subsequently assessed their potential effect on MSCs, which are a crucial part of the wound healing process. Analyzing general *in vitro* characteristics such as population doubling time and multi-lineage differentiation, we found different stimulating and inhibiting effects. Adipogenesis and osteogenesis are known antagonistic MSC differentiations. Continuous exposure to RJEVs strongly increased osteogenic differentiation, while decreasing adipogenic differentiation. Pre-conditioning MSCs with RJEVs had a similar tendency, however, the effect was not significant. Interestingly, chondrogenic differentiation was not influenced by pre-conditioning with RJEVs, but significantly increased when continuously exposed to RJEVs. These results indicate that no increase in the multipotency of MSCs can be observed, since not all three lineages are increased in a similar manner, as described for e.g. extracorporeal shockwave treatment in MSCs (43). Population doubling time increased over time in the control and RJEV pre-conditioned groups (after an initial decrease), which is a known effect in MSCs (44,45). RJEVs had a preserving effect on MSC proliferation over time, with no significant changes in PDT, and the potential underlying effects should be investigated in future studies.

Most interesting for wound healing is the secretory profile of MSCs, since tissue-resident MSCs modulate the wound bed by secreting a variety of proteins. Among the most important ones are i) IGF, a chemotactic agent for endothelial cells that stimulates hyaluronan as well as increasing fibroblast and keratinocyte migration (46); ii) FGF, that displays an anti-fibrotic effect by decreasing myofibroblast differentiation and fibronectin (47,48); iii) HGF, that increases migration, proliferation, and matrix metalloproteinase production of keratinocytes as well as increases dedifferentiation of epidermal cells (49–51), and iv) VEGF, that increases angiogenesis, collagen deposition, and epithelization (52). Overall, RJEVs significantly increased IGF, HGF, and VEGF secretion, but not FGF, indicating a more pro-angiogenic and pro-migratory effect in wound healing, and only minor effects on scarring and fibrosis.

Furthermore, the anti-inflammatory effect of Royal Jelly has previously been described in several studies (17,53,54), and has subsequently been attributed to three RJ-specific fatty acids (10-hydroxydecanoic acid (10-H2DA), Trans-10-hydroxy-2-decenoic acid (10-HDAA) and 1, 10-decanedioic acid (SEA)) that modulate the MAPK pathway (54). Interestingly, the observed inhibiting effects were minimal, compared to the fatty acid dose applied. The content of 10-H2DA in RJ is between 1.5-2.5% (55). Chen et al. utilized 2.5-5mM 10-H2DA, which corresponds to the equivalent of around 37 g of crude Royal Jelly, indicating a secondary mechanism of action for anti-inflammatory pathways. In our experiments (utilizing between 10^5^ to 10^7^ RJEVs, equivalent to 0.1-10 mg crude RJ), we found a significant reduction in pro-inflammatory cytokine secretion for IL1β, IL6, and TNFα in the higher concentration, and for IL6 and TNFα in the lower concentration (Fig. 4A). Furthermore, we could verify the involvement of RJEVs in the inhibition of JNK and ERK1/2 phosphorylation (Fig. 4B). The observed effects for RJEVs are comparable in magnitude and dosage to crude RJ, as reported by Kohno et al for the same cell line (RAW 264.7) (17), indicating a stronger involvement of RJEVs in the anti-inflammatory effect of RJ as honeybee specific fatty acids.

In literature, the *in vivo* effects of Royal Jelly in wound healing models are well-described and have been associated with an interplay of previously mentioned fatty acids, MRJP1, and antibacterial peptides such as Defensin-1 (56). We hypothesized that RJEVs act as protection against degradation and transport vehicles for such proteins or peptides. In our previous study, we reported type-I-collagen as a suitable delivery matrix for RJEVs, displaying a stable release of functional RJEVs for at least 7 days (22). Therefore, we tested RJEV antibacterial and pro-regenerative properties in separate animal models in order to comply with the 3R concept. Antibacterial assays were performed in a subcutaneous pocket, reducing the risk of systemic infection, and we observed strong bactericidal effects of RJEVs that were released from type-1 collagen gels. The observed results for RJEVs were comparable to previously reported *in vitro* effects for the same bacterial strain *S. aureus* ATCC 25923 (24), while untreated control groups displayed substantial bacterial growth within the surgical pocket (Fig. 5).

Regarding *in vivo* wound healing assays, RJEVs significantly accelerated initial wound closure compared to controls. However, no difference was observed at the end of the study (day 15) since all wounds were fully closed (Fig.6B). Similar effects were observed by Bucekova et al, utilizing honeybee defensin-1 (56), a peptide also identified within RJEVs (Fig.1C). The initial increase in tissue regeneration might be explained by RJEVs modulating the inflammatory phase, which is backed up by our prior *in vitro* findings associated to a significant increase in fibroblast and MSC migration (22,24), bactericidal effects (Fig. 5), modulation of the tissue-resident MSC secretome (Fig. 3), and modulation of LPS-stimulated immune-responses (Fig. 4). The advanced wound healing state can further be observed in the decrease of inflammatory cells and the increase of epithelial cells (Fig. 6C). While a shortening of the inflammatory phase has only minor effects on physiological wound healing, it plays a crucial role in chronic wounds that are prone to prolonged inflammation (57). Overall, RJEVs continuously delivered through a biomaterial matrix display promising effects as an anti-inflammatory treatment for chronic wounds, which have to be further investigated in future studies.

Summarizing, we have gained first insights into the biogenesis of RJEVs by confirming the presence of the exosomal markers CD63 and syntenin and found that RJEVs carry important active cargo proteins such as MRJP1, Defensin-1 and Jellein-3. RJEV uptake analysis revealed a potential double mechanism of action within the cell, with the direct release of cargo into the cytoplasm after membrane fusion, and intracellular sorting after macropinocytosis or clathrin-dependent endocytosis. Furthermore, RJEVs have demonstrated to modulate MSCs on several levels (e.g. differentiation, proliferation, secretome) as well as LPS-induced inflammation in macrophages. In *in vivo* studies, we confirmed the known anti-bacterial effects of RJEVs, and discovered an acceleration of wound healing in a splinted mouse model. Overall, Royal Jelly is a highly complex bee product that has been of interest for medical applications for decades. In the last years, several active compounds have been discovered and RJEVs appear to be the missing link elucidating some of the yet unexplained effects. Further studies are needed, not only for potential medical application of RJEVs, but also basic research on honeybee cell functions, in order to understand the biogenesis, uptake, and the roles of RJEVs for honeybees themselves.

## Acknowledgments

This work was supported by ANID FONDECYT Iniciación grant N°11180406, FONDECYT grant N°1220803 and FONDECYT grant N°1220804. Furthermore, we would like to thank Luis Délano for his help and support with the histological samples. Figures 5A, 6A, and 7 were created with BioRender.com.

